# Toll-like receptor 4 signaling in osteoblasts is required for load-induced bone formation in mice

**DOI:** 10.1101/2022.08.05.502963

**Authors:** Ibtesam Rajpar, Gaurav Kumar, Paolo Fortina, Ryan E. Tomlinson

## Abstract

During skeletal development, expression of nerve growth factor (NGF) leads to the survival of afferent sensory nerves that express neurotrophic tyrosine kinase receptor type 1 (TrkA), the high affinity receptor for NGF. In adulthood, NGF is expressed by mature osteoblasts following mechanical loading and signals through TrkA receptors in resident sensory nerves to support load-induced bone formation. However, the regulation of NGF in osteoblasts following loading is not well understood. In this study, we sought to determine whether osteoblastic expression of toll-like receptor 4 (TLR4), a key receptor in the NF-κB signaling pathway, is required to initiate NGF-TrkA signaling to support skeletal adaptation following mechanical loading. First, we observed that NF-κB inhibition reduces NGF expression induced by axial forelimb compression. Moreover, we observed that TLR4+ periosteal cells are increased after mechanical loading. Therefore, we generated a novel mouse model in which *Tlr4* is selectively removed from the mature osteoblast lineage. Although *Tlr4* conditional knockout mice have normal skeletal mass and strength in adulthood, the loss of TLR4 signaling results in significant reductions in periosteal lamellar bone formation following axial forelimb compression. Furthermore, we demonstrated that the upregulation of *Ngf* following application of fluid shear stress to calvarial osteoblasts is significantly reduced by NF-κB and TLR4 inhibitors. Finally, RNA sequencing demonstrated that the deficits in load-induced bone formation in CKO mice can be attributed to dysregulated inflammatory signaling. In total, our study reveals a novel role for TLR4 in skeletal adaptation to mechanical loading in bone, which may enable new therapeutic strategies for diseases of low bone mass and provide new targets for musculoskeletal pain relief.

## INTRODUCTION

The skeleton is constantly subjected to various compressive, tensile, and shear forces. The cellular constituents of bone, namely the osteoblasts and osteocytes, respond to these loads by depositing new bone at regions of high strain and removing bone at regions of low strain, in a process known as strain adaptive bone remodeling^1; 2^. Thus, remodeling of bone tissue is achieved by the conversion of mechanical cues to biochemical signals by bone cells^3^. One mechanism by which these signals are perceived is through the dense network of peripheral nerves present on the mammalian bone surface^4^. The vast majority of these nerves express the neurotrophic tyrosine kinase receptor type I (TrkA)^5; 6^, which is the high affinity receptor for the classic neurotrophin nerve growth factor (NGF)^7^. During development, NGF is produced by osteochondral progenitors as part of a developmental program to specify the amount and type of innervation required in bone^8^. In adulthood, NGF is also produced by mature osteoblasts in response to loading, and NGF-TrkA signaling is required to achieve maximal load-induced bone formation^9 10^. As a result, inhibition of TrkA signaling in adult mice significantly reduces bone formation following axial forelimb compression^10^. Furthermore, exogenous NGF injection in wild-type mice increases load-induced bone formation rate and Wnt signaling in osteocytes^10^. However, more investigation is needed to understand the control of NGF expression by osteoblasts involved in a well-orchestrated anabolic response to loading.

Previous studies have observed that NF-κB signaling is rapidly activated in osteoblastic cells by mechanical forces^11-13^. For example, application of fluid shear stress to osteoblasts results in the rapid increase of intracellular calcium, which is required for the release and nuclear translocation of NF-κB factors to activate the production of prostaglandins^13^. Generally, the NF-κB factors have been associated with homeostatic functions in bone, primarily catabolic bone resorption^14^, although a recent study showed that the constitutive activation of NF-κB signaling in osterix-expressing cells increased overall bone mass through enhanced basal and load-induced bone formation^15^. Importantly, NF-κB activation is specifically required for B-cell-derived and NP-cell-derived NGF expression^16; 17^ and activation of NF-κB factors promotes the NGF-mediated survival of sympathetic neurons^18^. In total, these studies strongly suggest a role for NF-κB signaling in the transcriptional activation of *Ngf* in mature osteoblasts following osteoanabolic mechanical loading.

Among potential cell-surface receptors associated with NF-κB signaling, the toll-like receptors (TLR) have been classically associated with the maintenance of innate immunity but are also expressed in a variety of cells in the musculoskeletal lineage including chondrocytes, synoviocytes, and osteoblasts^19^. Here, we focus on TLR4 – a type I transmembrane receptor with an extracellular domain comprising of 19-25 copies of the leucine-rich repeat (LRR) motif that recognizes highly conserved molecular patterns of evolutionary origin, such as the classic lipopolysaccharide (LPS) and high mobility group box protein 1 (HMGB1)^20^. In bone, TLR4 activation has been linked to the increased expression of RANKL and M-CSF^21^. These signaling factors are activated during osteoclastogenesis and promote bone resorption, but also increase secretion of the osteoanabolic Wnts (Wnt3a and Wnt5a) to promote osteogenic differentiation, proliferation, and angiogenesis^19; 22^. Analogous to the NF-κB signaling proteins, the TLR4 receptor is activated in response to mechanical stimulation^23^. In a previous study with healthy human disc cells, both a TLR2 and TLR4 agonist were found to increase NGF gene expression following treatment^17; 24^. Further, inhibition of NF-κB signaling with a small molecule inhibitor in disc cells silenced TLR2-driven NGF expression at both the gene and protein levels^17^. Therefore, we hypothesized that activation of TLR4 in osteoblasts drives NF-κB signaling to promote the NGF expression required for normal bone formation in response to mechanical loading.

To investigate this hypothesis, we developed a conditional knockout mouse line in which *Tlr4* was specifically removed from the mature osteoblast lineage using osteocalcin Cre-mediated recombination. We analyzed adult mice for skeletal phenotype prior to loading, and bone formation and changes in the bone transcriptome following axial forelimb compression. We further confirmed our results with an *in vitro* model of fluid shear stress application in which osteoblasts were treated with various NF-κB inhibitors and a TLR4-specific inhibitor to determine the requirement of NF-κB signaling factors and the TLR4 receptor in the NGF response to loading. Our findings reveal novel mechanistic insights to the NGF-TrkA signaling pathway in bone, and a new role of the TLRs in osteoanabolic loading.

## METHODS

### Mice

Tlr4^fl/fl^; Osteocalcin (OC)-Cre+ mice (CKO) on a C57BL/6J background were generated for this study in accordance with IACUC of Thomas Jefferson University. These mice were generated by crossing mice carrying Tlr4 homozygous floxed alleles (Jackson Labs #024872) with mice carrying Cre recombinase driven by the osteocalcin promoter (Jackson Labs #019509). Tlr4^fl/fl^ (WT) littermates were included as controls in the study. NGF-EGFP mice express EGFP under the control of the full length mouse *Ngf* promoter^25^. All mice were housed at 72 degrees and fed a daily diet of rodent chow (LabDiet 5001).

### Skeletal preparations

P0 mice were euthanized by cervical dislocation and placed in ice cold PBS. Immediately following euthanasia, pups were skinned and eviscerated. Following dehydration in ethanol and tissue permeabilization in acetone, skeletons were incubated for 3-4 days in a staining solution consisting of 0.3% alcian blue stock and 0.1% alizarin red S stock at 37°C. Excess stain was removed by soft tissue hydrolysis in 1% potassium hydroxide, which enabled transparency and visualization of skeletal elements. Skeletons were imaged using a dissecting microscope.

#### *In vivo* mechanical loading

Axial forelimb compression of the right forelimb was performed using a material testing system (TA Instrument Electroforce 3200 Series III) with custom designed fixtures. Prior to loading, mice were anesthetized with 3% isoflurane and buprenorphine (0.12mg/kg, IP), and they were maintained under 1.5% isoflurane for the duration of the loading protocol. Each mouse was loaded using a 2Hz sinusoidal, rest-inserted waveform with a peak force of 3N for 100 cycles each day over three consecutive days (D0-D2). This loading protocol has previously been shown to induce robust lamellar bone formation at the mid-shaft of the ulna^26^. The non-loaded left forelimb served as a contralateral control.

### Histomorphometry

To assess load-induced bone formation by dynamic histomorphometry, adult (16-20 weeks) male and female mice were given calcein (10mg/kg, Sigma C0875) on D3 and alizarin red s (30mg/kg, Sigma A3882) on D8 of the loading protocol by IP injection. Animals were euthanized on D10. Forelimbs were dissected, fixed in 10% formalin overnight at 4°C, dehydrated in 70% ethanol and embedded in polymethylmethacrylate. 100 micron thick sections from the mid-diaphysis of the ulnae were cut using a precision saw (Isomet 1000) and mounted on glass slides with Eukitt mounting medium (Sigma 03989) and visualized by confocal microscopy (Zeiss). Bone formation parameters, including endosteal (Es) and periosteal (Ps) mineralizing surface (MS/BS), mineral apposition rate (MAR), and bone formation rate (BFR/BS), were quantified using ImageJ according to the guidelines of the ASBMR Committee for Histomorphometry Nomenclature^27^.

To characterize bone cells by static histomorphometry, 8-9 week old femurs were fixed, decalcified in 14% EDTA for 2 weeks at 4°C, and embedded in paraffin. 4 micron thick sections were stained with either standard H&E or tartrate-resistant acid phosphatase (TRAP) stain. Sections were imaged using a bright field microscope (EVOS M5000 imaging system). Three 20X regions of trabecular bone were analyzed to determine average number of osteoblasts or osteoclasts per bone surface and osteocytes per bone area.

For immunofluorescent staining of TLR4 protein, adult mice were given three bouts of compressive loading, and forelimbs were harvested on D7 post loading on D0. Forelimbs were fixed, decalcified and embedded in paraffin, and 4 micron thick sections were cut from the mid-section of the ulnar diaphysis. Sections were deparaffinized, rehydrated and treated with hot, 0.05% citrate buffer for 30 minutes. Next, non-specific antigens were blocked with a goat serum, before overnight incubation with a rabbit, anti-mouse TLR4 antibody (Abcam) at 4°C. The next day, sections were washed and treated with a goat anti-rabbit IgG fluorescent secondary antibody (Alexa Fluor™ 488, Life Technologies) for 1 hour. Sections were treated with a nuclear counterstain (Vectashield mounting medium, Vector Labs, Newark, CA) and imaged with a confocal microscope (Zeiss). Three high-powered 10X regions of the periosteum were analyzed to determine average number of TLR4 positive cells as a percentage of total periosteal cells.

To visualize NGF-EGFP expression, forelimbs from NGF-EGFP mice were harvested 3 hours following one bout of compressive loading, fixed, decalcified, and embedded in O.C.T compound (Tissue-Tek), and 10 micron thick sections were cut from the mid-section of the ulnar diaphysis. Sections were mounted with media containing DAPI (Vectashield H1500) and imaged with a confocal microscope (Zeiss).

### Micro computed tomography and three-point bending

For analysis of skeletal phenotype by microCT, the right femurs were dissected post-euthanasia from adult mice and frozen at -20°C in PBS-soaked gauze. Each bone was scanned using a Bruker microCT analyzer fixed with a 1mm aluminum filter, and with scanning parameters of 55 kV and 181 uA at a resolution of 12 microns. Bone scans were reconstructed using nRecon (Bruker), and aligned and analyzed using CTan (Bruker) for cortical and trabecular bone parameters. For mechanical testing of adult femurs with three-point bending, femurs were placed on custom fixtures with the condyles facing down and a span length of 7.6 mm measured with calipers. Femurs were then fractured to failure with a monotonic displacement ramp of 0.1 mm/s. Force and displacement data from the test as well as geometric parameters from microCT were analyzed using a custom GNU Octave script to derive structural and material properties.

### RNA isolation and sequencing

The middle third of loaded and non-loaded ulnae were harvested 3 hours after a single bout of loading on D0 to extract RNA for whole transcriptome analysis. Bones were centrifuged to remove bone marrow, and stored in RNAlater (Thermo Fisher) at -80 °C. Bone tissue was pulverized in TRIzol (Thermo Fisher) to obtain uniform tissue homogenates, and total RNA was isolated from bone homogenates using the TRIzol method as per manufacturer’s instructions. RNA was further purified using a Qiagen RNeasy kit (Qiagen) and samples with RIN values ranging from 5-8 were selected for sequencing. Illumina Stranded Total RNA libraries were prepared including Ribo-Zero Plus rRNA depletion as per the manufacturer’s protocols. Libraries were sequenced on the Illumina NovaSeq 6000 using 2×100 base paired-end chemistry v1.5 reagents. The resulting reads in FASTQ format were aligned to the mouse genome version GRCm38 with Gencode M19 transcript annotations using the RSEM-STAR pipeline^28; 29^. Gene expression count files produced by RSEM were analyzed for differential expression using the DESeq2 package in R/Bioconductor software^30^. The Gene Set Enrichment Analysis (GSEA) pre-ranked tool was applied to genes ranked by their resulting DESeq2 test statistic to identify differentially regulated Hallmark genesets^31; 32^. Additional visualization was performed using the R/Bioconductor EnhancedVolcano package (version 1.8.0).

### Cell lines

The MC3T3-E1 Subclone 4 pre-osteoblast cell line was derived from ATCC^®^ (#CRL-2593^™^). Primary osteoblasts were isolated from D0-D3 CKO and WT mice using standard techniques^33^. Briefly, neonatal calvaria were dissected, cut into pieces, washed in sterile PBS, and sequentially digested using 1.8 mg/ml type I collagenase (Worthington Biochemical Corp. #4197) to release the osteoblasts into solution. For loading experiments, cells were seeded in standard 96-well plates and cultured in complete media comprising α-Minimal Essential Medium (Corning #15-012-CV), 10% fetal bovine serum (Corning #35-010-CV) and 1% penicillin/streptomycin antibiotic (Sigma #P4333).

#### *In vitro* mechanical loading and NFkB pathway inhibitors

To assess *Ngf* expression, MC3T3-E1 cell cultures were subjected to fluid shear stress of 1 ml/min for 0 (control), 10, 15, 30, and 60 minutes using a previously developed microfluidics fluid flow device^34^ coupled to a standard peristaltic pump (Fisher) with 1.6 mm diameter polyethylene tubing. To determine the requirement of the NFkB pathway following loading, osteoblast cultures were supplemented with either 5 uM BAY 11-7082 (Sigma) or 50 uM pyrrolidine dithiocarbamate (PDTC, Sigma), 1 hour before loading for 60 minutes at 1ml/min. To determine the requirement of TLR4 signaling for *Ngf* expression, cultures were supplemented with either 5 or 10 uM TAK-242 (Cayman Chemicals), 2 hours before loading for 60 minutes at 1ml/min. Cells were harvested 1 hour following loading in all inhibitor experiments. Control groups were not loaded (WT) and/or treated with DMSO.

### Gene expression analysis

The expression of bone markers *in vitro* was determined using real time quantitative PCR. Total RNA from cell cultures was isolated using the TRIzol method, purified with the Qiagen RNeasy kit, and quantified using a NanoDrop spectrophotometer. RNA was reverse transcribed using iScript cDNA synthesis kit (BioRad), and amplified using SYBR Green biochemistry with the QuantStudio 3 real time PCR system (Applied Biosystems). Relative gene expression was quantified using the ΔΔCT method, using GAPDH as a reference gene. Primer sequences were designed using Primer-BLAST and purchased from IDT Technologies. A complete list of primer sequences is provided in Table 1.

**Table 1.**
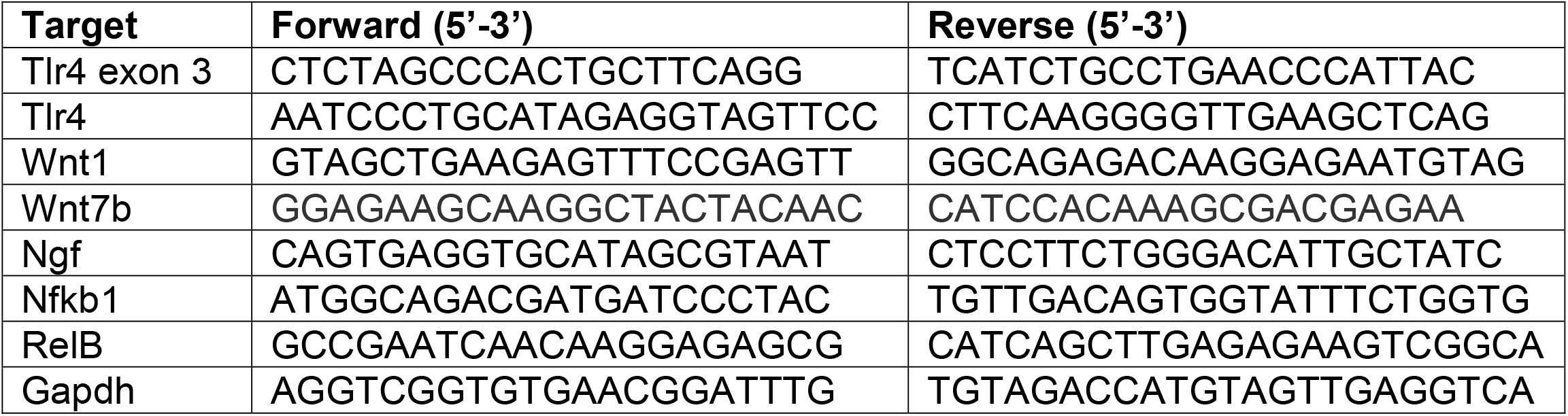
Oligonucleotide primers used for qRT-PCR.

### Statistics

Data was collected in a blinded fashion from all animals in the study, unless removed prior to analysis due to illness or injury after consultation with veterinarian staff. After data collection, outlier analysis was performed using Grubbs’ test if necessary. For *in vivo* comparisons of bone formation parameters in loaded versus non-loaded bone, two-way ANOVA was used with post hoc Tukey’s tests. For analysis of skeletal phenotype, two-tailed, unpaired t-tests were used. Multiple group means for *in vitro* experiments were compared using one-way ANOVA with post hoc Tukey’s tests (p ≤ 0.05). Statistical analysis was carried out in Prism 9 (Graphpad).

## RESULTS

### Loading increases TLR4+ periosteal cells as well as NGF expression that is blocked by NF-_κ_B inhibition

To determine the requirement of NF-κB signaling for NGF expression in bone following axial forelimb compression, we administered the NF-κB inhibitor BAY 11-7082 (20 mg/kg, Cayman Chemicals) or vehicle (DMSO) to adult NGF-EGFP mice 1 hour before a single bout of axial forelimb compression. By confocal microscopy, we observed robust NGF expression at the periosteal surface 3 hours after loading that was essentially silenced by the inhibitor (Fig. 1A,B). Next, mRNA was harvested 24 hours after loading for analysis by qRT-PCR. Here, we found that treatment with BAY 11-7082 was associated with significant decreases in *Ngf* (−52%), *Wnt1* (−37%), and *Wnt7b* (−46%), such that they were not different than control (non-loaded) limbs (Fig. 1C-E). In addition, *Nfkb1* was significantly decreased (−44%) in mice treated with BAY 11-7082 (Fig. 1F). Since NF-κB signaling can be activated through toll-like receptors, including TLR4, we examined the expression of TLR4 in loaded and non-loaded forelimbs using immunohistochemistry 7 days after axial forelimb compression. Here, we observed that the number of TLR4+ periosteal cells was significantly increased in loaded limbs (Fig. 1H-I). In total, these data suggested that NF-κB signaling following loading induced NGF expression and was potentiated through the activity of TLR4. To test this hypothesis, we generated TLR4 conditional knockout (CKO) mice in which exon 3 of *Tlr4* was excised by Cre-mediated recombination under the control of the full-length osteocalcin promoter.

**Figure 1.**
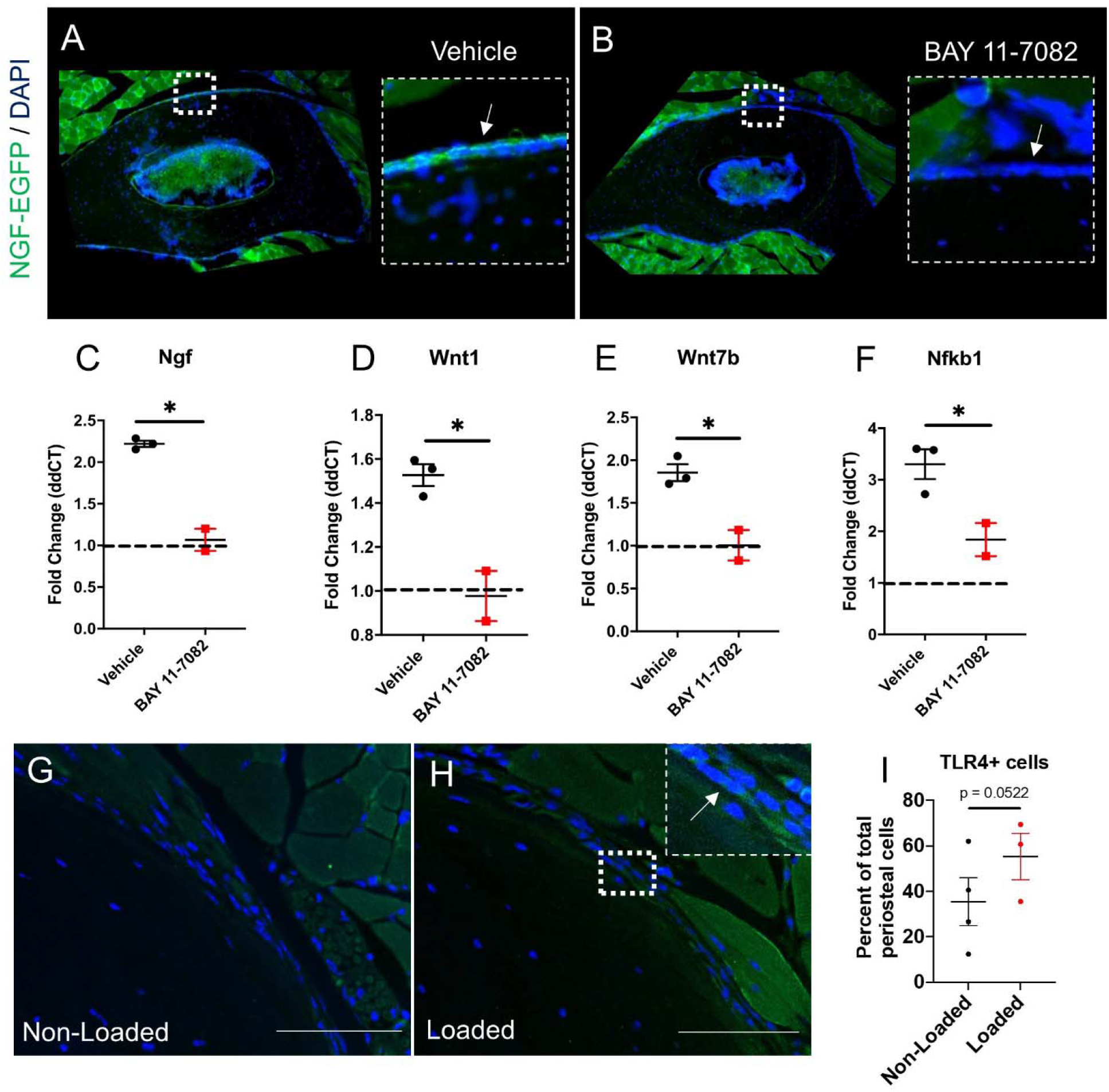
Inhibition of NF-_κ_B decreases NGF and Wnt signaling in loaded bone. A) Expression of NGF in the periosteum of loaded ulna from NGF-EGFP mice was reduced by B) BAY 11-7082 3 hours after loading. Similarly, C) NGF, D) Wnt1, E) Wnt7b, and F) Nfkb1 expression were decreased 24 hours after loading. G) Non-loaded and H) loaded ulnar sections stained for TLR4 7 days after loading, with arrow indicating TLR4+ cells. I) Quantification of TLR4+ cells in both loaded and non-loaded sections as a percentage of total periosteal cells. Scale bars are 100 microns. * p < 0.05 vs. vehicle. n=2-4 per group.

### Adult bone mass and strength is not affected by loss of TLR4 in the mature osteoblast lineage

TLR4 CKO mice appeared healthy and fertile and were obtained at the expected Mendelian frequency. The genotype of the mice was confirmed by amplification of genomic DNA as well as *Tlr4* mRNA. Alizarin red and alcian blue skeletal preparations of neonates at postnatal day 0 revealed no significant differences in the size or shape of skeletal elements (Supp. Fig. 1). Paraffin sections of adolescent (8-9 weeks) femurs were analyzed to assess changes in cellular composition of CKO mice. While no changes were noted in numbers of osteoblasts or osteocytes, TRAP staining revealed a significantly reduced number of osteoclasts (−25%) in CKO metaphyseal bone as compared to WT littermates (Supp. Fig. 2). Next, we analyzed adult bone mass and geometry using femurs harvested from 16-20 week old mice. MicroCT revealed no significant differences in trabecular or cortical bone parameters in either male or female WT and CKO mice (Supp. Fig. 3, 4). Similarly, femoral three-point bending did not reveal any significant differences in ultimate stress, Young’s modulus, or ultimate bending energy in CKO femurs compared to WT (Supp. Fig. 5). In total, constitutive loss of TLR4 in the mature osteoblast lineage did not significantly affect osteoblasts, osteocytes, bone mass, or bone strength during development or in adulthood.

### Osteoblastic TLR4 is required for load-induced bone formation

To directly test our hypothesis that signaling through TLR4 contributes to the osteoanabolic response following mechanical loading, male and female CKO mice were subjected to three consecutive days of axial forelimb compression. Forelimbs labeled with calcein and alizarin red were harvested 10 days after loading for analysis by dynamic histomorphometry. Here, we observed robust lamellar bone formation on the periosteal and endosteal bone surfaces of the ulnar mid-diaphysis in loaded limbs harvested from WT mice (Fig. 2A, I), including significant increases in mineralizing surface per bone surface (MS/BS), mineral apposition rate (MAR), and bone formation rate per bone surface (BFR/BS) in loaded limbs compared to non-loaded contralateral control limbs (Table 2). However, the loss of TLR4 greatly diminished the anabolic response, with relative (loaded – non-loaded) periosteal BFR/BS significantly reduced (−50%) in male CKO mice following loading (Fig. 2E). In female mice, we observed similar response to loading in WT forelimbs, but female CKO mice had even larger reductions in relative periosteal (−68%) and endosteal (−47%) BFR/BS (Fig. 2M, P), which was mainly driven by decreases in MAR (−67%) as compared to WT littermates (Fig. 2L). In non-loaded limbs, bone formation parameters were not significantly different between genotypes (Table 2).

**Figure 2.**
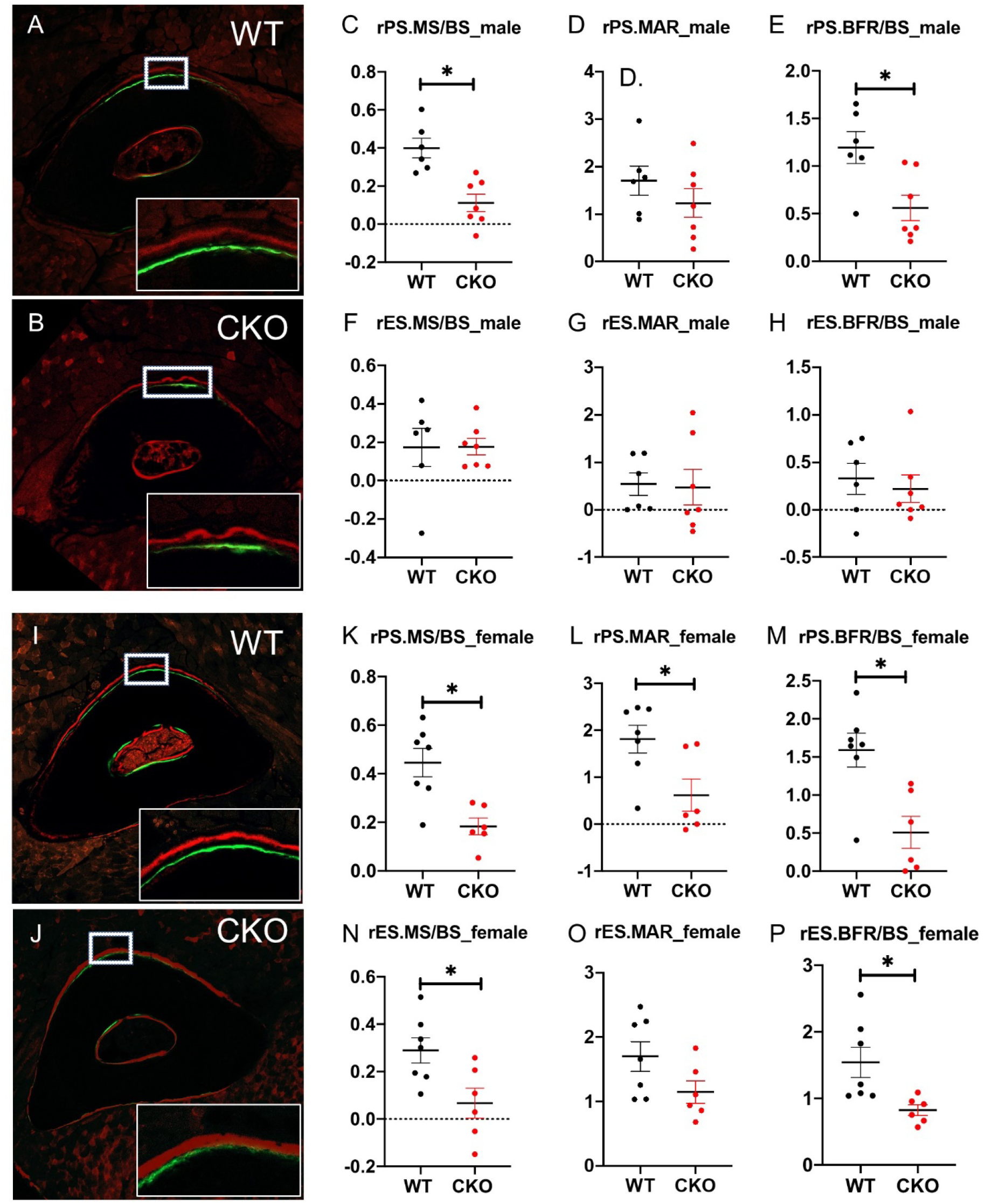
Tlr4 is required for lamellar bone formation following compressive forelimb loading. Impaired bone formation following loading in *Tlr4* conditional knockout mice is demonstrated by representative cross-sections from the ulnar mid-diaphysis of A) WT and B) CKO male mice with quantification of D-F) periosteal and F-H) endosteal parameters. Similar results were obtained in females, as demonstrated by representative cross-sections from the ulnar mid-diaphysis of I) WT and J) CKO female mice with quantification of K-M) periosteal and N-P) endosteal parameters. * p < 0.05 between genotype. n=6-7 per group.

**Table 2.** Loss of TLR4 diminished load-induced bone formation in both female and male mice by dynamic histomorphometry. Values are presented as mean ± standard deviation. * p < 0.01 vs. non-loaded, # p < 0.01 vs. CKO by two-way ANOVA.

| Female |  |  |  |  |  |  |
| --- | --- | --- | --- | --- | --- | --- |
| CKO | PS.MS/BS<br>($\mu\text{m}/\mu\text{m}$) | Es.MS/BS<br>($\mu\text{m}/\mu\text{m}$) | Ps.MAR<br>($\mu\text{m}/\text{day}$) | Es.MAR<br>($\mu\text{m}/\text{day}$) | Ps.BFR/BS<br>($\mu\text{m}^3/\mu\text{m}^2/\text{d}$<br>ay) | Es.BFR/BS<br>($\mu\text{m}^3/\mu\text{m}^2/\text{d}$<br>ay) |
| Loaded | 0.46 ± 0.06 | 0.56 ± 0.06 | 1.58 ± 0.37 | 2.16 ± 0.2* | 0.86 ± 0.22 | 1.38 ± 0.17* |
| Non-Loaded | 0.28 ± 0.07 | 0.49 ± 0.07 | 0.96 ± 0.25 | 1.02 ± 0.24 | 0.35 ± 0.07 | 0.56 ± 0.14 |
| WT | PS.MS/BS<br>($\mu\text{m}/\mu\text{m}$) | Es.MS/BS<br>($\mu\text{m}/\mu\text{m}$) | Ps.MAR<br>($\mu\text{m}/\text{day}$) | Es.MAR<br>($\mu\text{m}/\text{day}$) | Ps.BFR/BS<br>($\mu\text{m}^3/\mu\text{m}^2/\text{d}$<br>ay) | Es.BFR/BS<br>($\mu\text{m}^3/\mu\text{m}^2/\text{d}$<br>ay) |
| Loaded | 0.69 ± 0.03 | 0.75 ± 0.03 | 2.78 ±<br>0.22*,# | 2.42 ±<br>0.18*,# | 1.93 ±<br>0.19*,# | 1.85 ±<br>0.21*,# |
| Non-Loaded | 0.24 ± 0.04 | 0.46 ± 0.03 | 0.97 ± 0.28 | 0.72 ± 0.15# | 0.34 ± 0.10 | 0.31 ± 0.05# |
| Male |  |  |  |  |  |  |
| CKO | PS.MS/BS<br>($\mu\text{m}/\mu\text{m}$) | Es.MS/BS<br>($\mu\text{m}/\mu\text{m}$) | Ps.MAR<br>($\mu\text{m}/\text{day}$) | Es.MAR<br>($\mu\text{m}/\text{day}$) | Ps.BFR/BS<br>($\mu\text{m}^3/\mu\text{m}^2/\text{d}$<br>ay) | Es.BFR/BS<br>($\mu\text{m}^3/\mu\text{m}^2/\text{d}$<br>ay) |
| Loaded | 0.42 ± 0.03 | 0.31 ± 0.05 | 1.99 ± 0.34* | 0.79 ± 0.36 | 0.81 ± 0.15 | 0.32 ± 0.14 |
| Non-Loaded | 0.31 ± 0.03 | 0.12 ± 0.02 | 0.76 ± 0.15 | 0.31 ± 0.07 | 0.25 ± 0.04 | 0.09 ± 0.01 |
| WT | PS.MS/BS<br>($\mu\text{m}/\mu\text{m}$) | Es.MS/BS<br>($\mu\text{m}/\mu\text{m}$) | Ps.MAR<br>($\mu\text{m}/\text{day}$) | Es.MAR<br>($\mu\text{m}/\text{day}$) | Ps.BFR/BS<br>($\mu\text{m}^3/\mu\text{m}^2/\text{d}$<br>ay) | Es.BFR/BS<br>($\mu\text{m}^3/\mu\text{m}^2/\text{d}$<br>ay) |
| Loaded | 0.57 ± 0.01 | 0.44 ± 0.09 | 2.46 ± 0.34* | 1.03 ± 0.21# | 1.38 ±<br>0.18*,# | 0.57 ± 0.13 |
| Non-Loaded | 0.17 ± 0.04 | 0.26 ± 0.07 | 0.75 ± 0.34 | 0.49 ± 0.15 | 0.18 ± 0.04 | 0.24 ± 0.07 |

### Loss of TLR4 signaling in osteoblasts reduced *Ngf* expression following fluid shear stress

Next, we assessed if activation of TLR4 signaling induced *Ngf* expression in osteoblasts by utilizing an *in vitro* model of fluid shear stress. First, we utilized MC3T3-E1 cells to determine the time course of expression of *Ngf* following the application of fluid shear stress. Here, we found incremental increases in *Ngf* transcription in response to increased duration of fluid shear stress. Specifically, *Ngf* transcription was significantly increased following 30 minutes (+1.5 fold) and 60 minutes (+2.0 fold) of fluid shear stress as compared to baseline (Fig. 3B). Next, we assessed mRNA expression in calvarial osteoblasts harvested from CKO neonates and WT littermates to determine baseline gene expression. Consistent with our overall hypothesis, we observed significant downregulation in the expression of *Tlr4* (−40%), *Ngf* (−50%), and *Nfkb1* (−40%) in CKO osteoblasts as compared to WT (Fig. 3C). To assess the requirement of the NF-κB pathway in *Ngf* expression following application of fluid shear stress, culture media of primary calvarial osteoblasts was supplemented with the NF-κB inhibitors BAY 11-7082 or PDTC^35; 36^. Both inhibitors diminished the expression of *Ngf* (−25%) in calvarial osteoblasts at 1 hour following 60 minutes of loading (Fig. 3D), although *Nfkb1* expression was not affected (Fig. 3F). Finally, to delineate the specific role of the NF-κB pathway receptor TLR4, both MC3T3-E1 cells and calvarial osteoblasts from CKO mice were treated with TAK-242, a small molecule TLR4 inhibitor^37^. In this experiment, *Ngf* expression was significantly increased (+92%) following fluid shear stress and significantly reduced by inhibitor in a dose-dependent manner. Specifically, *Ngf* transcription was decreased –33% with 5uM (p = 0.0484) and –67% with 10uM (p = 0.0045) of TAK-242 (Fig. 3G). We also observed that *Ngf* transcription was also reduced in CKO osteoblasts treated with 10uM TAK prior to loading (Fig. 3H). In total, these data demonstrate that inhibition of TLR4 activity impairs *Ngf* expression in osteoblasts following fluid shear stress.

**Figure 3.**
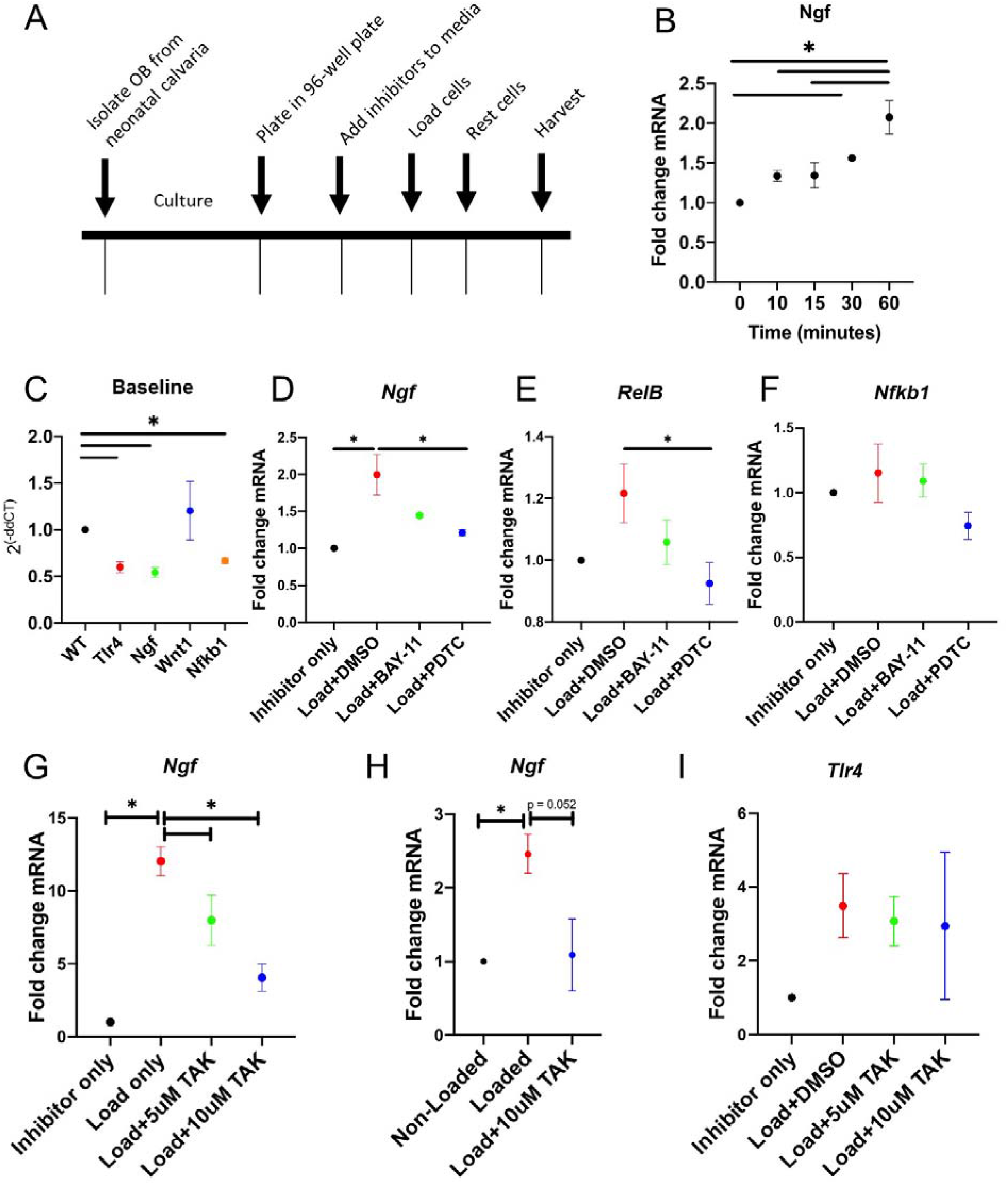
TLR4 signaling is required for upregulation of *Ngf* following fluid shear stress in osteoblasts. A) Timeline illustrates the study design. B) Expression of *Ngf* 0-60 minutes after loading in MC3T3 cells. C) Gene expression of *Tlr4, Ngf, Wnt1, and Nfkb1* in CKO osteoblasts as compared to wildtype (WT) at baseline (not loaded). D,E) Load-induced upregulation of *Ngf* and *RelB* was significantly reduced by pretreatment with BAY11-7082 and/or PDTC prior to loading. F) Expression of *Nfkb1* was not affected by loading or inhibition. G) Load-induced upregulation of *Ngf* in MC3T3 cells was significantly reduced by TAK-242 in a dose-dependent manner. H) Similarly, load-induced upregulation of *Ngf* in primary osteoblasts was significantly reduced by TAK-242. I) *Tlr4* expression was not affected by inhibition of signaling. * p < 0.05 by one-way ANOVA.

### Loss of *Tlr4* in bone dysregulates inflammatory signaling following loading

To further characterize the role of osteoblastic TLR4 in osteogenic mechanical loading, mRNA was isolated from the middle third of loaded and non-loaded ulnae harvested from both CKO and WT littermates 3 hours after a single bout of axial forelimb compression. RNA sequencing revealed 200 transcripts that were differently expressed (fold change > 2 and an adjusted p-value of ≤ 0.05) between loaded and non-loaded bones of wildtype and CKO mice (Supplemental Table 1). Principal component analysis (PCA) revealed a distinct clustering of loaded and non-loaded bone, with PC1 accounting for 30.1% and PC2 accounting for 21.5% of the total variance (Fig. 4A). Several differentially expressed transcripts were significantly upregulated in loaded bone when compared to non-loaded, including known mechanoresponsive genes such as *FosB* (+8.0X), *Wnt1* (+10.3X) and *Ngf* (+5.7X) (Fig. 4C). In *Tlr4* CKO mice, osteoanabolic genes such as *Dmp1* (+3.1X), *Bmp4* (+2.5X), and *Bmp7* (+2.1X) were also significantly upregulated in loaded limbs. Gene set enrichment analysis was performed to identify the relevant biological pathways responsive to loading (Supplemental Table 2). Here, we observed significant upregulation of WNT (p = 0.008) and TGFβ (p = 0.000) signaling following loading in both genotypes of mice (Fig. 4D). In contrast, several inflammatory pathways, including IL2 (p = 0.026), IL6 (p = 0.000), IFN gamma (p = 0.000), and IFN alpha (p = 0.000), were upregulated in WT mice but downregulated in CKO mice following loading (Fig. 6A, 6B). Surprisingly, the loss of Tlr4 in CKO mice appears to result in the activation of signaling pathways not directly related to bone accrual following loading, such as myogenesis and adipogenesis (Fig. 6A).

**Figure 4.**
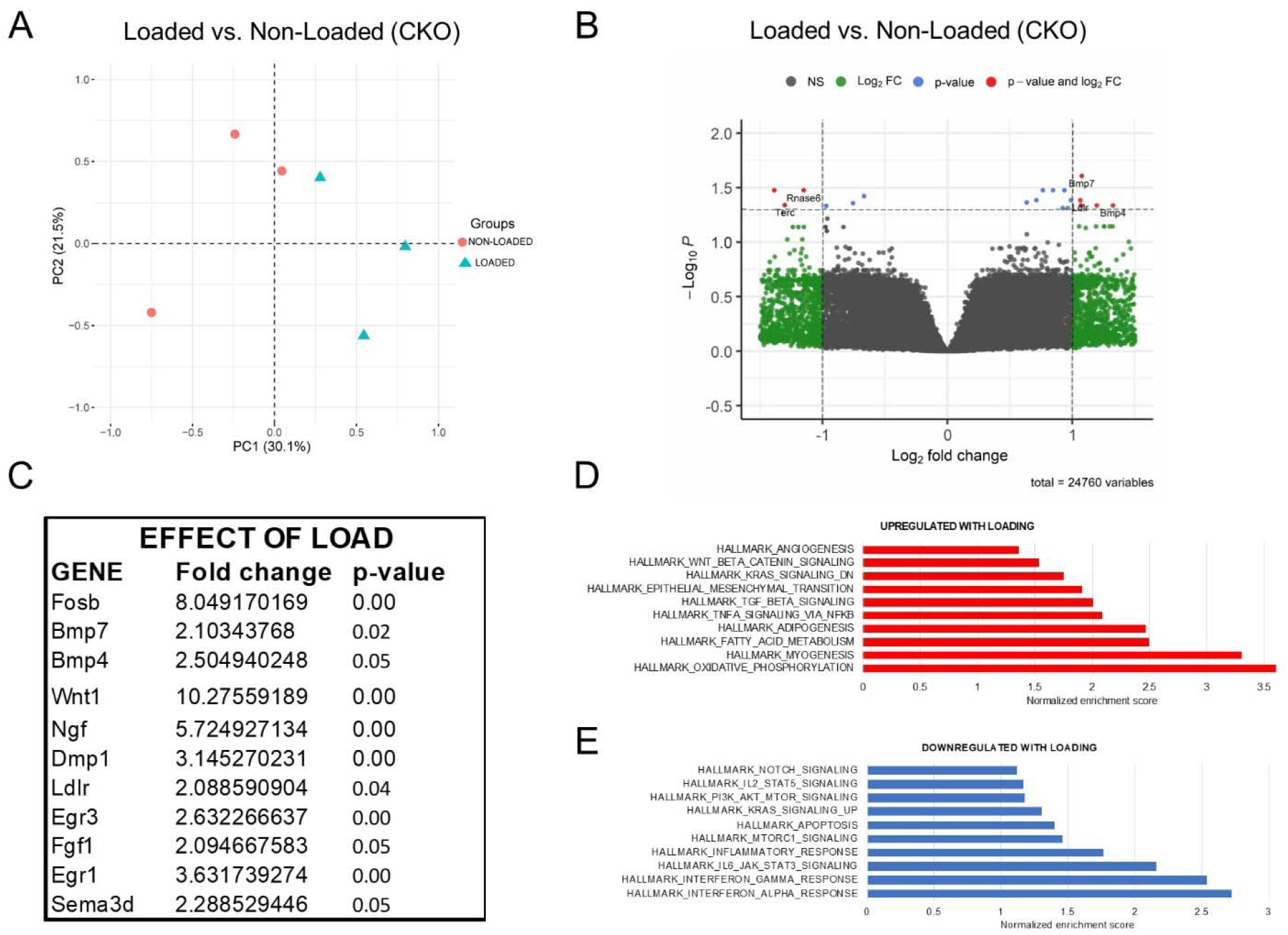
Loading induced significant changes in the bone transcriptome of CKO mice. RNA sequencing was performed on loaded and non-loaded forelimbs from CKO adult mice harvested 3 hours following loading. A) Principal component analysis of loaded vs. non-loaded bones. B) Volcano plot of differentially expressed transcripts following loading. C) Subset of differentially expressed genes relevant to anabolic response to loading. Relevant signaling pathways that were significantly D) upregulated or E) downregulated following loading. n = 4-7 per group.

PCA also revealed the distinct clustering of transcripts isolated from CKO and WT non-loaded bones, with two principal components accounting for 39.5% of the total variance (Fig. 5A). Here, we observed 161 differentially expressed gene transcripts, with 34 significantly downregulated and 127 significantly upregulated in non-loaded bones from CKO as compared to WT (Fig. 5B). Similar to the response to loading, we noted that several gene transcripts related to inflammatory signaling were aberrantly upregulated in the non-loaded limb of CKO mice, such as the inflammatory mediators *Il5ra* (+2.4X, p = 0.0003), *Hmgb3* (+2.1X, p = 0.0004), and *Tnf* (+2.0X, p = 0.0004) (Fig. 5C). Furthermore, gene set enrichment analysis was used to identify numerous inflammatory processes, such as MTORC and TNFα signaling, that were significantly upregulated in CKO bones (Fig. 5D). Indeed, these signaling pathways were upregulated in both loaded and non-loaded conditions in CKO mice (Fig. 6C). Furthermore, the loss of Tlr4 in CKO mice also resulted in a downregulation of a variety of signaling pathways in non-loaded conditions, including TGFβ signaling and angiogenesis (Fig. 6D). In total, these data demonstrate that TLR4 signaling in the osteoblast is necessary for proper inflammatory signaling in both loaded and non-loaded bones.

**Figure 5.**
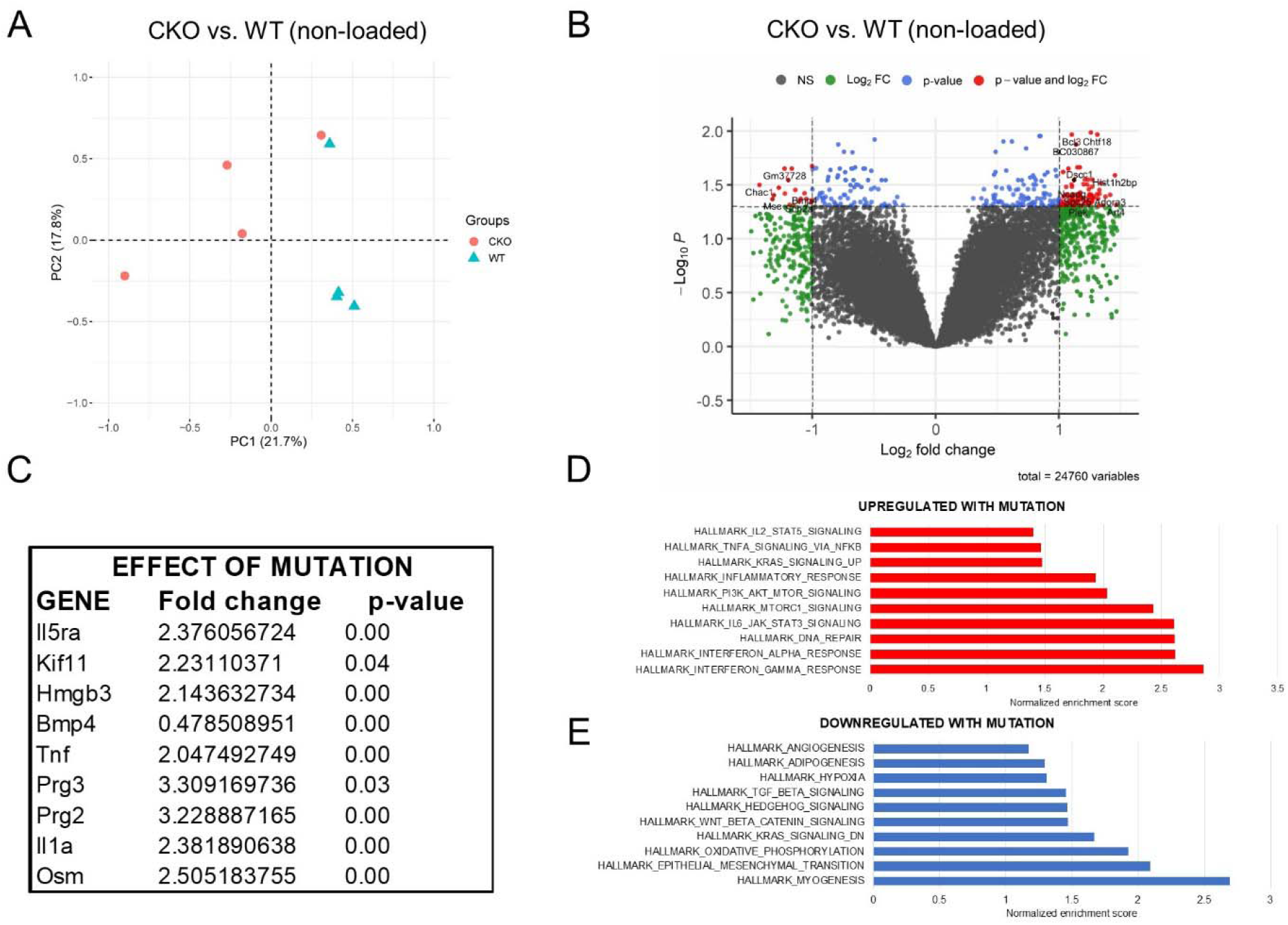
Loss of *Tlr4* induced significant changes in the bone transcriptome. RNA sequencing was performed on wildtype and CKO non-loaded forelimbs. A) Principal component analysis of CKO vs. WT bones. B) Volcano plot of differentially expressed transcripts. C) Subset of differentially expressed genes relevant to the mutation. Relevant signaling pathways that were significantly D) upregulated or E) downregulated in CKO vs. wildtype. n = 4-7 per group.

**Figure 6.**
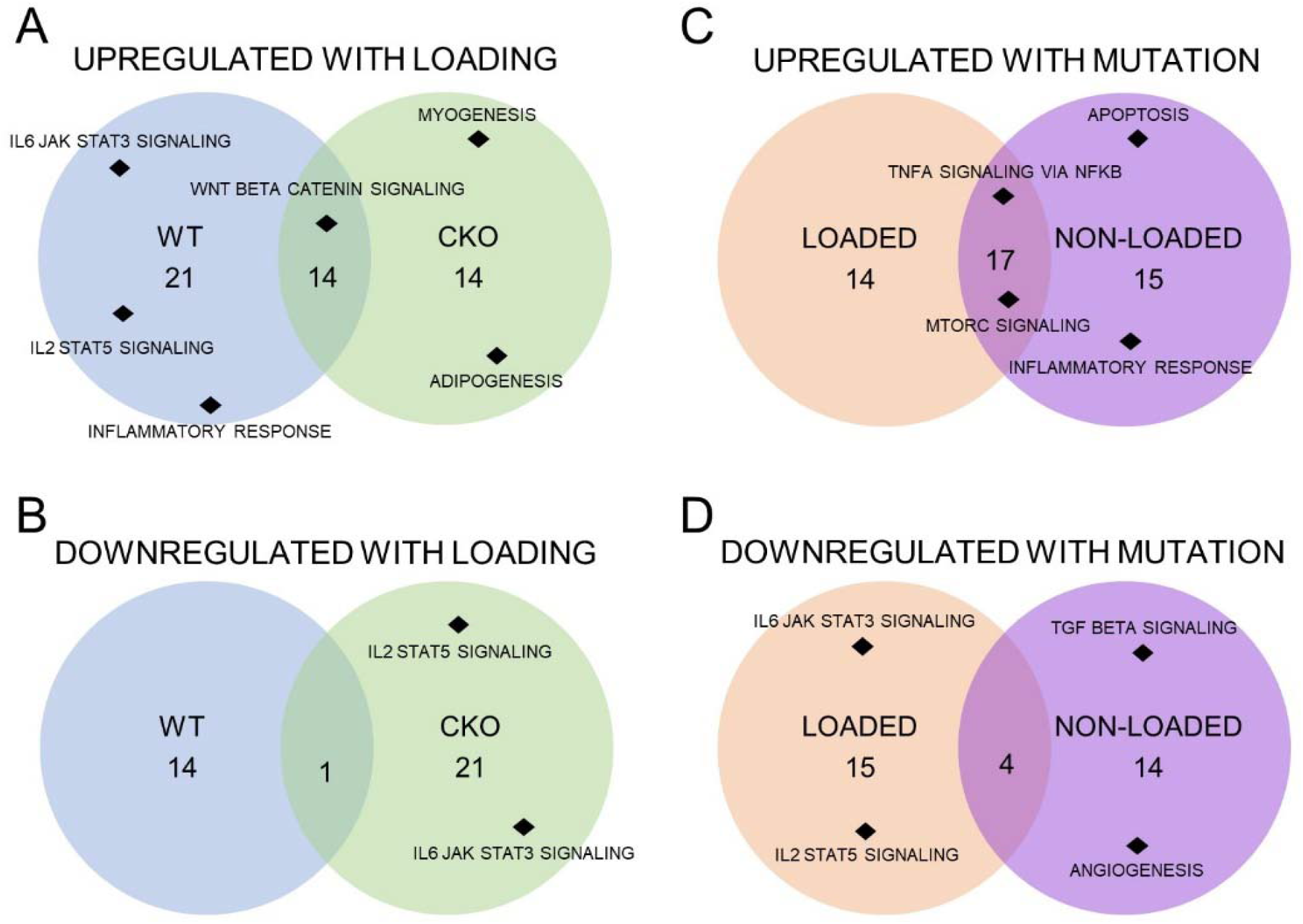
Common and unique hallmarks from RNA sequencing. The number of signaling pathways A) upregulated or B) downregulated in WT mice, CKO mice, or both genotypes in loaded limbs as compared to non-loaded limbs. Similarly, the number of pathways C) upregulated or D) downregulated in loaded limbs, non-loaded limbs, or both limbs in CKO mice as compared to WT mice. A full list of the significant pathways is available in Supplemental Table 2. n = 4-7 per group.

## DISCUSSION

In this study, we determined the requirement and specific role of osteoblastic TLR4 signaling following axial forelimb compression in mice. Specifically, we observed that loss of *Tlr4* in the osteoblast lineage did not significantly affect the size or shape of the skeleton and had minimal effects on cellular composition. However, CKO mice formed significantly less bone in response to axial forelimb compression as compared to WT littermates, and had a dysregulated inflammatory response as demonstrated by RNA sequencing. Furthermore, we showed *in vitro* that *Ngf* expression in response to fluid shear stress was impaired in a dose-dependent manner by TLR4 inhibitors. In total, these data strongly support a model in which activation of TLR4 on mature osteoblasts induces the expression of NGF through NF-κB signaling to support load-induced bone formation (Fig. 7).

**Figure 7.**
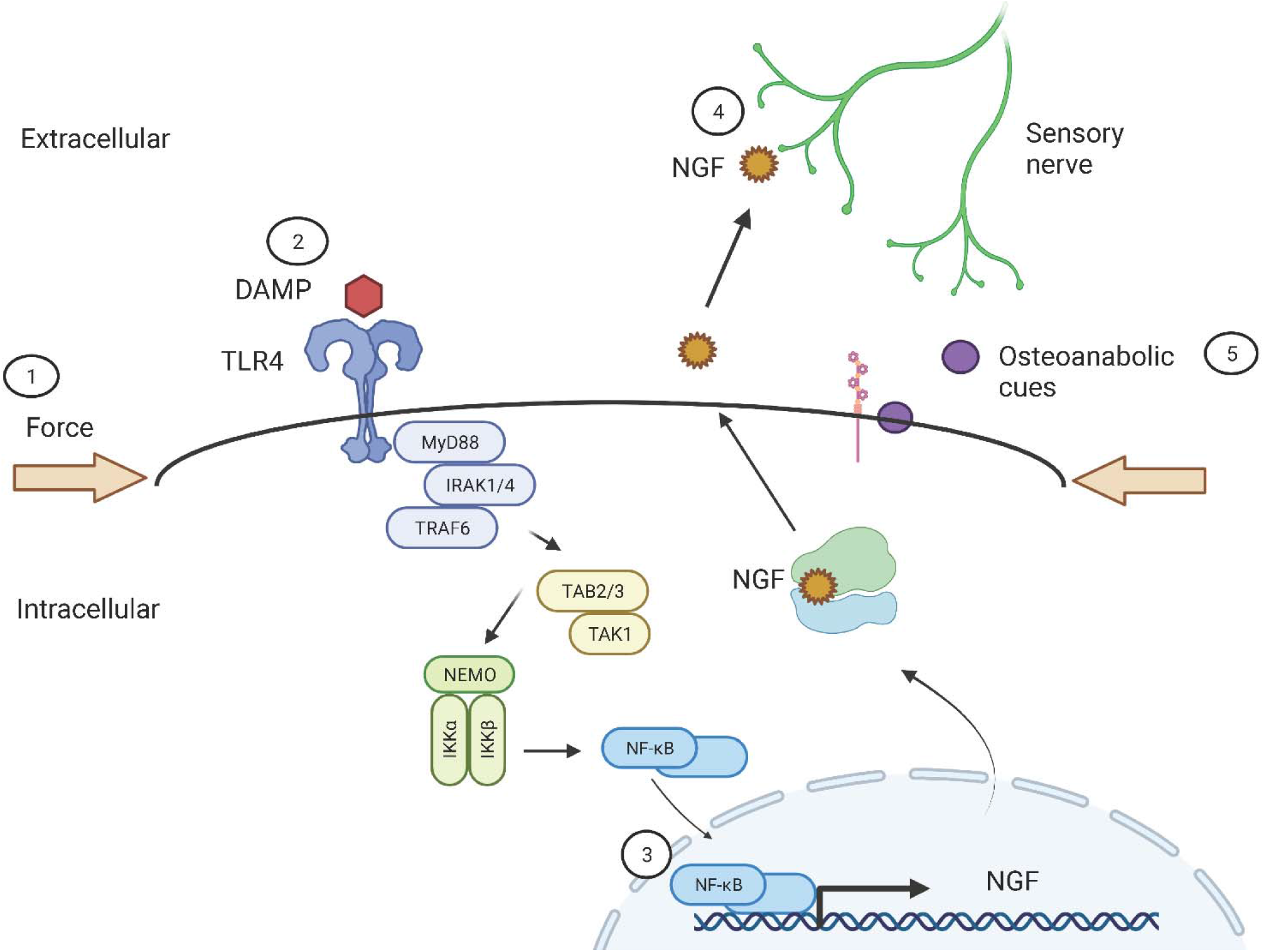
Schematic of TLR4 signaling in load-induced bone formation. 1) Mechanical loading of osteoblasts promotes the activation of TLR4 receptors by endogenous DAMPS and/or TLR4 agonists, 2) TLR4 activation stimulates formation of the NEMO complex, resulting in the release and nuclear translocation of NF-κB factors, 3) NF-κB factors bind and activate transcription of *Ngf*, 4) NGF is released from osteoblasts and binds its high affinity receptor TrkA on skeletal sensory nerves, and 5) NGF-TrkA signaling promotes the release of osteoanabolic cues that support new bone formation.

To pinpoint the function of TLR4 in osteoblasts during load-induced bone formation, we developed a novel conditional knockout mouse model of *Tlr4* deficiency using osteocalcin Cre-mediated deletion to target the mature osteoblast lineage. We hypothesized that the relatively late expression of osteocalcin in mature osteoblasts would circumvent potential developmental deficits due to loss of *Tlr4*. Consistent with this hypothesis, mutant CKO mice did not exhibit any differences in bone parameters as determined by microCT or material properties by standard three-point bending when compared to their WT littermates. This finding is consistent with global and myeloid-cell specific *Tlr4* knockout models that did not report any skeletal abnormalities^38^. However, loss of *Tlr4* has been shown to impact the morphology and function of the retina, cardiac function, and developmental neuroplasticity in mice^39^. Nonetheless, we did observe lower numbers of osteoclasts in the trabecular bone of CKO mice (S. Fig. 2E-G). This finding did not appear to affect bone mass or outcomes from mechanical loading and likely stems from the altered inflammatory signaling due to loss of osteoblastic TLR4 (Fig. 5). Nonetheless, the reduced osteoclast number may be relevant to a future studies of TLR4 function in bone, particularly regarding callus remodeling following fracture. Indeed, previous studies have demonstrated that TLR4 activation is associated with bone resorption and osteoblast-mediated osteoclastogenesis during active bone remodeling and in inflammatory bone disorders^19; 40^.

Furthermore, our results demonstrate that activation of TLR4 in mature osteoblasts is required for load-induced bone formation, illustrating a novel osteoanabolic function of a receptor primarily associated with inflammatory bone loss^38^. In particular, we observed increased numbers of TLR4+ cells on the ulnar periosteal surface following axial forelimb compression. More study is required to determine if these TLR4-expressing cells generated after loading are mature osteoblasts, osteoprogenitors, or non-osteoblastic cells present in periosteum^41^. To investigate sex-related differences in the response of bone to *Tlr4* knockout, both male and female mice were included in our analysis. Loss of *Tlr4* in both male and female mice resulted in similar overall reductions in the key outcome parameter following axial forelimb compression – periosteal bone formation rate (BFR/BS). This result was mainly driven by decreased activation of osteoblasts, as demonstrated by significantly decreased mineralizing surface per bone surface (MS/BS) in both male and female mice; only female mice also had significantly decreased mineral apposition rate (MAR). Furthermore, only female mice displayed a significant reduction in endosteal BFR/BS, which was also mainly driven by decreased endosteal MS/BS. In total, it appears that loss of osteoblastic *Tlr4* affected female mice more strongly than male mice, but more study is required to determine the source of this difference. Furthermore, there were no significant differences in bone formation parameters between genotypes in non-loaded limbs, underscoring that the role of TLR4 appears limited to the response to loading.

We specifically hypothesized that TLR4 signaling may be required for *Ngf* expression through NF-κB signaling in osteoblasts following mechanical loading. First, we showed that administration of NF-κB inhibitor to NGF-EGFP reporter mice silences NGF expression following axial forelimb compression. Next, we utilized a series of *in vitro* studies to show that *Ngf* expression following the application of fluid shear is significantly reduced by both BAY11-7082 and PDTC, inhibitors of the NF-κB signaling pathway, as well as TAK-242, a TLR4-specific inhibitor, in a dose-dependent manner. As a result, we were surprised to find that both wildtype and CKO mice displayed a significant upregulation of *Ngf* following axial forelimb compression by RNA sequencing (Supplemental Table 1). More study is required to determine if the source of this NGF is outside of the mature osteoblast lineage (e.g. inflammatory cells or osteoprogenitors) or if constitutive loss of *Tlr4* results in the expression of alternative receptors that can compensate for the loss of TLR4 in mature osteoblasts. Nonetheless, CKO mice did not activate several inflammatory signaling pathways observed to be upregulated in WT mice following loading, including IL2, IL6, and IFNα signaling. Thus, loss of osteoblastic *Tlr4* results in a dysregulated inflammatory response, suggesting that TLRs act as regulators of the inflammation that potentiates load-induced bone mass accrual^42^. Furthermore, previous studies have illustrated that activation of NGF-TrkA signaling in monocytes decreases TLR-mediated translocation of NF-κB factors and glycogen synthase kinase 3 activity, resulting in a decreased production of inflammatory cytokines^43; 44^. Therefore, additional studies are required to determine if TLR4-mediated *Ngf* transcription is involved in the proper resolution of inflammation following osteogenic mechanical loading.

In total, the results from this study illustrate the novel role of TLR4 signaling in mature osteoblasts to support load-induced bone formation. Furthermore, we showed that the TLR4-NGF-TrkA signaling axis in mature osteoblasts is dispensable for skeletal development but activated in response to mechanical loads to support bone formation and mediate inflammatory signaling. Future work will focus on the differentiation stage of osteoblasts that express TLR4, the specific receptor agonists required for load-induced bone formation, and the role of osteoblastic TLR4 in other skeletal contexts, such as fracture repair and osteomyelitis.

## ACKNOWLEDGEMENTS

Our research is supported by the National Institute of Arthritis and Musculoskeletal and Skin Diseases, the National Institute of Dental and Craniofacial Research, and the Office of the Director of the National Institutes of Health under award numbers AR074953 (RET), DE028397 (RET), and OD025128 (PF). The authors thank Dr. Michael Kawaja for supplying mice used in this study and Dr. Adam Ertel for help with data visualization. The content is solely the responsibility of the authors and does not necessarily represent the official views of the funding bodies. The authors have no conflicts of interest to disclose.

## DATA AVALIBILITY STATEMENT

Raw and processed RNA sequencing data for this study has been uploaded to the NCBI Gene Expression Omnibus (GEO) with the accession number GSE210597. Additional data that support the findings of this study are available from the corresponding author upon request.

## AUTHOR CONTRIBUTIONS

**Ibtesam Rajpar**: Conceptualization, Methodology, Investigation, Writing; **Gaurav Kumar:** Investigation; **Paolo M. Fortina**: Methodology, Supervision; **Ryan E. Tomlinson**: Conceptualization, Methodology, Investigation, Supervision, Funding acquisition, Writing. The authors have no conflicts of interest.

