## Supplemental Figures 1-5 for "Toll-like receptor 4 signaling in osteoblasts is required for load-induced bone formation in mice"

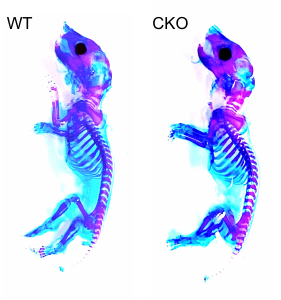

**Supplemental Figure 1.** **Skeletal preparation of postnatal D0 mice.** No differences were noted in the size or shape of skeletal elements in WT and CKO mice.

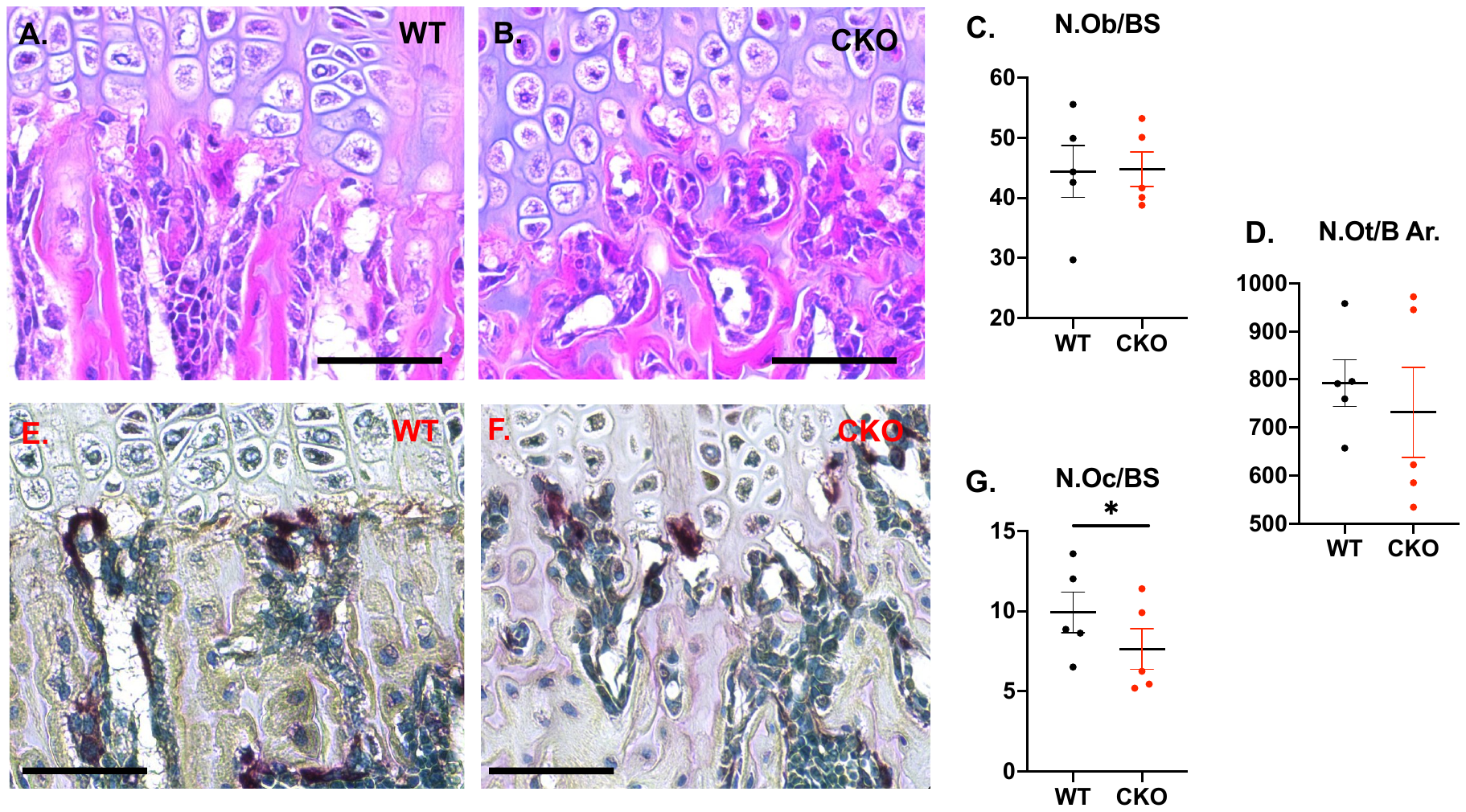

**Supplemental Figure 2.** **Quantification of bone cells in young mice.** Long sections of femurs from WT and CKO mice were stained with either standard H&E stain (A,B) to quantify osteoblasts per bone surface (C) and osteocytes per bone area (D), or TRAP stain (E,F) to quantify osteoclasts per bone surface (G) in trabecular bone. Scale bars are 50 microns. * denote significant differences by t-test (p ≤ 0.05).

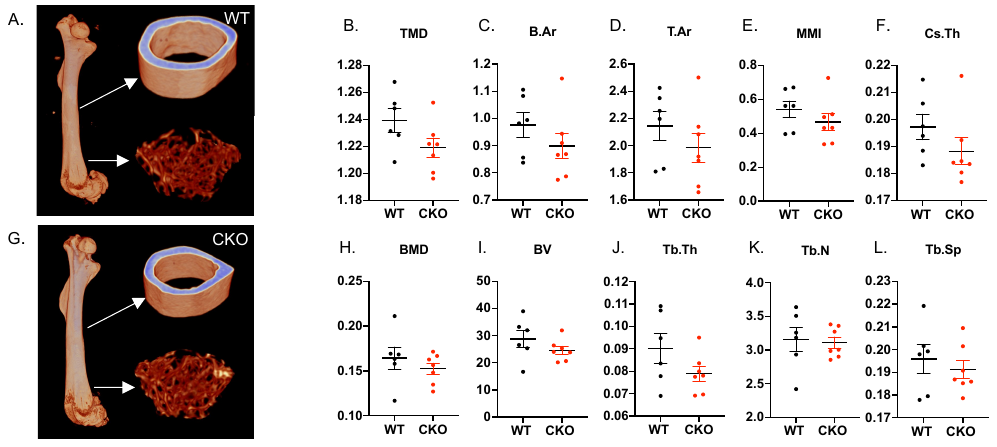

**Supplemental Figure 3.** **Quantification of cortical and trabecular bone parameters in adult male femurs.** White arrows in reconstructed images (A,G) indicate ROI for cortical and trabecular bone in WT (A) and CKO (G) femurs. No differences were noted in cortical (B-F) or trabecular (H-L) bone parameters between WT and CKO mice.

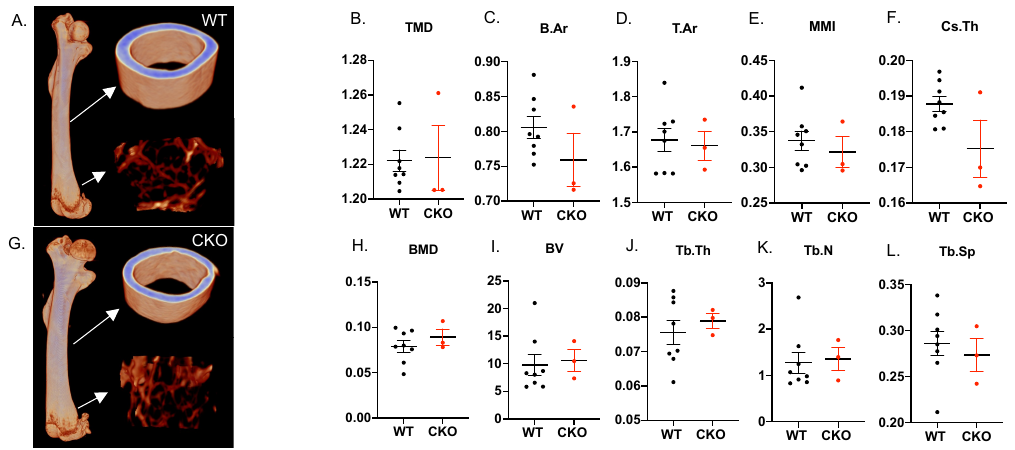

**Supplemental Figure 4. Quantification of cortical and trabecular bone parameters in adult female femurs.** White arrows in reconstructed images indicate ROI for cortical and trabecular bone in WT (A) and CKO (G) femurs. No differences were noted in cortical (B-F) or trabecular (H-L) bone parameters between WT and CKO mice.

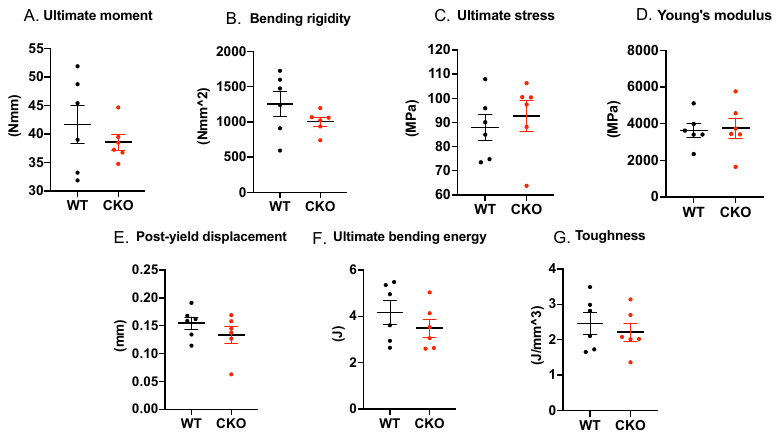

**Supplemental Figure 5. Mechanical testing of adult mouse femurs.** Femurs from WT and CKO mice were subjected to standard three-point bending and data were analyzed using a custom GNU Octave script to derive structural and material properties.
